# Resiliency to Alzheimer’s disease neuropathology can be distinguished from dementia using cortical astrogliosis imaging

**DOI:** 10.1101/2024.05.06.592719

**Authors:** Stephanie Barsoum, Caitlin S. Latimer, Amber L. Nolan, Alexander Barrett, Koping Chang, Juan Troncoso, C. Dirk Keene, Dan Benjamini

## Abstract

Despite the presence of significant Alzheimer’s disease (AD) pathology, characterized by amyloid β (Aβ) plaques and phosphorylated tau (pTau) tangles, some cognitively normal elderly individuals do not inevitably develop dementia. These findings give rise to the notion of cognitive ‘resilience’, suggesting maintained cognitive function despite the presence of AD neuropathology, highlighting the influence of factors beyond classical pathology. Cortical astroglial inflammation, a ubiquitous feature of symptomatic AD, shows a strong correlation with cognitive impairment severity, potentially contributing to the diversity of clinical presentations. However, noninvasively imaging neuroinflammation, particularly astrogliosis, using MRI remains a significant challenge. Here we sought to address this challenge and to leverage multidimensional (MD) MRI, a powerful approach that combines relaxation with diffusion MR contrasts, to map cortical astrogliosis in the human brain by accessing sub-voxel information. Our goal was to test whether MD-MRI can map astroglial pathology in the cerebral cortex, and if so, whether it can distinguish cognitive resiliency from dementia in the presence of hallmark AD neuropathological changes. We adopted a multimodal approach by integrating histological and MRI analyses using human postmortem brain samples. *Ex vivo* cerebral cortical tissue specimens derived from three groups comprised of non-demented individuals with significant AD pathology postmortem, individuals with both AD pathology and dementia, and non-demented individuals with minimal AD pathology postmortem as controls, underwent MRI at 7 T. We acquired and processed MD-MRI, diffusion tensor, and quantitative T_1_ and T_2_ MRI data, followed by histopathological processing on slices from the same tissue. By carefully co-registering MRI and microscopy data, we performed quantitative multimodal analyses, leveraging targeted immunostaining to assess MD-MRI sensitivity and specificity towards Aβ, pTau, and glial fibrillary acidic protein (GFAP), a marker for astrogliosis. Our findings reveal a distinct MD-MRI signature of cortical astrogliosis, enabling the creation of predictive maps for cognitive resilience amid AD neuropathological changes. Multiple linear regression linked histological values to MRI changes, revealing that the MD-MRI cortical astrogliosis biomarker was significantly associated with GFAP burden (standardized β=0.658, pFDR<0.0001), but not with Aβ (standardized β=0.009, *p*_FDR_=0.913) or pTau (standardized β=-0.196, *p*_FDR_=0.051). Conversely, none of the conventional MRI parameters showed significant associations with GFAP burden in the cortex. While the extent to which pathological glial activation contributes to neuronal damage and cognitive impairment in AD is uncertain, developing a noninvasive imaging method to see its affects holds promise from a mechanistic perspective and as a potential predictor of cognitive outcomes.

## Introduction

Alzheimer’s Disease (AD) is the most common form of dementia of older adults, and 6.7 million Americans aged 65 and older are living with AD as of 2023.^1^ The disease is characterized by initial memory loss and eventually impairments in executive functioning, spatial reasoning, and behavioral changes and dementia as it progresses.^2,3^ Future projections suggest that the prevalence of AD will increase as the general population continues to age,^4^ emphasizing the need for advancing the development of early diagnostic technologies with increased accuracy.

The neuropathologic hallmarks of AD, amyloid β (Aβ) plaques and neurofibrillary tangles (NFTs) of hyperphosphorylated tau (pTau), generally correlate with a clinical history of cognitive impairment or dementia.^5–7^ However, a number of clinical-pathologic studies reveal that cognitive impairment does not uniformly follow the presence of amyloid plaques and NFTs.^8–11^ Supporting these observations, about 30% of cognitively healthy elderly individuals exhibit amyloid positivity,^10,11^ and 13% show pTau positivity^12^ on PET scans. Complementing these findings, about 7% of cognitively intact older brain donor participants in aging and dementia studies have high pTau neuropathology, and up to 45% have intermediate levels of post-mortem pTau burden.^13^ Notably, examination of brains from individuals aged 85 and older consistently show some degree of AD neuropathologic change in the absence of dementia, exceeding expected prevalence rates.^14–16^ Following these observations, the concept of ‘resilience’ (or ‘asymptomatic AD’) emerged, signifying the absence of cognitive decline despite a burden of AD neuropathology that is normally associated with impaired cognition.^17–20^ These insights suggest that mechanisms beyond Aβ and pTau pathology play pivotal roles in driving tissue injury responses, leading to brain functional changes and cognitive impairment in AD.

The role of neuroinflammation and particularly of glial cells in the pathophysiology of AD has gained significant interest, driven by discoveries of genomic risk loci in genes associated with the human immune system.^21^ Microglial and astrocytic responses are increased in cortical regions in donor brains with high pTau burden,^22^ and glial cortical responses are evident early in the disease and intensify in proximity to amyloid plaques and NFTs as AD symptoms progress clinically.^23,24^ While the extent to which pathological glial activation contributes to neuronal damage and cognitive impairment in AD is uncertain, developing a noninvasive imaging method to see its affects holds promise from a mechanistic perspective and as a potential predictor of cognitive outcomes.

Development of noninvasive imaging techniques, especially those based on magnetic resonance imaging (MRI), to image neuroinflammation, and in particular, astrogliosis, has been challenging. This difficulty primarily arises from the limitations of conventional MRI methods in detecting astrogliosis, and in disentangling the response to cellular alterations induced by astrogliosis from the responses to co-morbid concurrent microstructural and chemical processes. Biophysical models applied to diffusion MRI data have been recently suggested to capture neuroinflammation. These were shown to correlate with histological measures in animal models but not in humans.^25^ Model driven diffusion MRI findings were also correlated with CSF levels of Aβ_1-40_, Aβ_1-42_, and pTau in humans, but not with neuroinflammatory markers.^26^ Additionally, while such biophysical models offer advantages in specificity and interpretability, they rely on strict assumptions and incorporate fixed parameters to reduce fitting ambiguities,^27^ which may not be valid in disease states or even in heterogeneous brain regions.^28–30^

Multidimensional MRI (MD-MRI) is a term used for an alternative approach that simultaneously encodes diffusion and relaxation, replacing traditional voxel-averaged scalar values with distributions of quantitative metrics, free of a biophysical model.^31,32^ These distributions offer insights into the individual diffusion and relaxation properties of distinct water populations, along with their correlations. In addition to proving invaluable for investigating tissue microstructure^33–37^ and pathology,^38–41^ MD-MRI was recently used to map traumatic brain injury induced astrogliosis in white matter via a radiological-pathological multimodal study in human postmortem brain samples.^42^

The present study was designed to test whether MD-MRI can provide maps of astroglial pathology in the cerebral cortex, and if so, can it distinguish cognitive resiliency from dementia in the presence of hallmark AD neuropathological changes. To confirm or refute these hypotheses, we have taken a multimodal approach integrating histology and MRI and conducted a detailed study on human postmortem brain samples from (a) non-demented individuals with minimal AD pathology postmortem; (b) non-demented individuals with significant AD pathology postmortem (referred to as “resilient”); and (c) individuals with both AD pathology and dementia. Careful co-registration of MRI and microscopy data allows quantitative multimodal analyses, in which the targeted specificity of immunostaining is utilized to determine sensitivity and specificity of MD-MRI towards Aβ, pTau, and astrogliosis. We show that cortical astrogliosis has a characteristic MD-MRI signature, which can be used to create maps that predict cognitive resilience in the presence of AD neuropathological changes. We believe that this framework, capable of discerning individuals with dementia from those with preserved cognition, holds promise in pinpointing those at risk of developing clinical dementia symptoms among asymptomatic elderly individuals positive for amyloid and pTau, thereby informing the development of innovative therapies aimed at averting cognitive decline in the presence of significant burden of AD neuropathology.

## Materials and Methods

### Donor specimens

We evaluated 13 brains donated for research from two different human brain collections. Five cases were obtained from the University of Washington (UWA) BioRepository and Integrated Neuropathology (BRaIN) laboratory. The use of human subject material was performed in accordance with the Declaration of Helsinki and the guidelines set by the UWA Institutional Review Board, including a waiver of informed consent. Consent for brain donation was obtained from the donor or from the legal next of kin, according to the protocols approved by the UW Institutional Review Board (IRB). The remaining eight donor brains were obtained from Johns Hopkins University (JHU), in which eight cases were participants in the Baltimore Longitudinal study on Aging (BLSA), and one donor was a participant in the Alzheimer’s Disease Research Center (ADRC) whose tissues were procured through the Johns Hopkins Brain Resource Center (BRC). Both the tissue donation consent process and the accessioning of tissues by the BRC are conducted under protocols approved by the JHU IRB.

The procurement, preservation, and preparation protocols for research donor brain tissues for this study were generally aligned across the two biorepositories, with some minor differences. Briefly, upon removal, one hemisphere undergoes fixation in 10% neutral-buffered formalin for at least two weeks while the contralateral hemibrain may be rapidly sliced and frozen depending on postmortem interval, or the whole brain is fixed. All the fixed brains are sliced coronally at 4 mm (UWA) or 10 mm (JHU) intervals, photographed, and sampled according to NIA-AA guidelines supplemented with biorepository-specific strategies that include sampling any gross lesions. Sampled tissues are processed, sectioned, stained, and evaluated according to National Institute of Aging-Alzheimer’s Association (NIA-AA) guidelines for AD neuropathologic change and AD-related disorders.^43,44^

For the present study, de-identified formalin-fixed tissue blocks (not processed or paraffin-embedded) of approximately 20 x 20 x 10 mm^3^ were dissected from cerebral cortical regions of the cases identified above. The specimens were sent to the National Institute on Aging (NIA) specifically for the current study and subsequently scanned using MD-MRI acquisitions. Upon completion, the specimens were returned and histologically processed as described above. A detailed description of donor demographics and characteristics is listed in Table 1.

**Table 1.** Main demographic and histopathological findings.

| Case | Age | Sex | Cognitive state | Braak | CERAD | Classification | Region |
| --- | --- | --- | --- | --- | --- | --- | --- |
| 1 | 91 | F | No dementia | III | Absent | Control | SFG |
| 2 | 30 | M | No dementia | 0 | Absent | Control | SFG |
| 3 | 64 | M | No dementia | I | Absent | Control | SFG |
| 4 | 65 | F | No dementia | I | Absent | Control | SFG |
| 5 | 83 | F | No dementia | III | Absent | Control | Temporal pole |
| 6 | 92 | F | No dementia | V | Moderate | Resilient | SFG |
| 7 | 71 | M | No dementia | IV | Frequent | Resilient | SFG |
| 8 | 89 | F | No dementia | V | Frequent | Resilient | SFG |
| 9 | 94 | M | Dementia | VI | Frequent | AD dementia | SFG |
| 10 | 91 | M | Dementia | VI | Frequent | AD dementia | Middle frontal cortex |
| 11 | 90 | F | Dementia | VI | Frequent | AD dementia | SFG |
| 12 | 91 | F | Dementia | VI | Frequent | AD dementia | SFG |
| 13 | 62 | F | Dementia | VI | Frequent | AD dementia | SFG |

Superior frontal gyrus (SFG).

### Definition of Study Groups

We studied three groups of donors: those with high AD neuropathology and dementia, those without significant AD neuropathology and cognitively intact, and those with high AD neuropathological changes but cognitively unimpaired. The AD subjects (referred to here as AD dementia) were donors who had received a clinical diagnosis of dementia according to the Diagnostic and Statistical Manual of Mental Disorders criteria^45^ and of probable AD according to the National Institute of Neurological Diseases and Stroke–Alzheimer’s Disease and Related Disorders Association criteria.^46^ The neuropathologic examination of these subjects showed neuritic Aβ plaque density with CERAD score of frequent and neurofibrillary tangle distribution of Braak stage VI. The brains were free of other age-related pathologies normally associated with dementia (see table 1). Control donors had no cognitive or behavioral deficits at the most recent cognitive evaluation, cerebrovascular disease, or history of alcohol/drug abuse. Neuropathological evaluation of these donors revealed CERAD cortical neuritic plaque score of none and Braak stage <IV (see table 1). Non-demented subjects with significant AD neuropathology (referred to here as resilient) exhibited no behavioral deficits, cerebrovascular disease, or alcohol/drug abuse history, and maintained cognitive integrity up to their last evaluation within 1-2 years before death. Neuropathological assessment revealed CERAD neuritic plaque density of moderate or frequent and Braak stage IV-V.

### MRI acquisition

Upon receipt at NIA and prior to MRI scanning, each formalin-fixed brain specimen was transferred to a phosphate-buffered saline filled container for 12 days to ensure that any residual fixative was removed from the tissue. Each specimen was then placed in a 20 mm tube, and immersed in perfluoropolyether (Fomblin LC/8, Solvay Solexis, Italy), a proton free fluid void of a proton-MRI signal. Specimens were imaged using a 7T Bruker vertical bore MRI scanner equipped with a microimaging probe and a 20 mm quadrupole RF coil.

Multidimensional diffusion-T_2_ data were acquired using a 3D diffusion-weighted sequence with a repetition time of 1000 ms and isotropic resolution of 200 µm. We used a previously published hierarchical sampling scheme^47^ to encode the multidimensional MR space spanned by T_2_ and mean diffusivity (i.e., T_2_-MD). Briefly, T_2_ was encoded by varying the echo time between 12.3 ms and 125 ms, and diffusion was encoded by varying the diffusion weighting b-value between 2400 s/mm^2^ and 14400 s/mm^2^, at 13 independent directions,^48^ with gradient duration of 4 ms and diffusion time of 15 ms. The total number of imaging volumes to generate voxelwise T_2_-MD distributions was 322. Full details regarding the MD-MRI acquisition can be found in the Supplementary Material.

A high-resolution MRI scan with an isotropic voxel dimensions of 100 µm was acquired using a fast low angle shot (FLASH) sequence^49^ with a flip angle of 49.6° to serve as a high resolution reference image and facilitate co-registration of histological and MR images.

A standard diffusion tensor imaging (DTI) protocol was applied with the same imaging parameters as the multidimensional data and using 21 diffusion gradient directions and four b-values ranging from 0 to 1400 s/mm^2^.

### Histology and Immunohistochemistry

After MRI scanning, each tissue block was transferred for histopathological processing. Tissue blocks from each brain specimen were processed using an automated tissue processor (details in the Supplementary Material). After tissue processing, each tissue block was embedded in paraffin, cut in a series of consecutive sections, and immunostained for hyperphosphorylated pTau (AT8) protein, for astrocytes (GFAP), for Aβ (6E10 or 4G8), for microglia (IBA1), and for pathological phosphorylated TDP-43 (pTDP-43). At least two sections per antibody and per sample were stained 1000 μm apart from each other, in accordance with the MRI slice thickness. More details regarding immunohistochemistry can be found in the Supplementary Material.

All stained sections were digitally scanned using either an Aperio whole slide scanner (Leica Biosystems, Richmond, IL) (UWA cases) or a Zeiss Axio Scan.Z1 digital slide scanner (Zeiss, Oberkochen, Germany) (JHU cases) at x20 magnification for further assessment and analyses. The scanned images underwent rigorous quality control measures to confirm positivity in control slides and identify staining artifacts or other features that might impede digital analysis.

### Quantification of histopathology

The following steps were taken to allow for a quantitative analysis of the microscopy data. First, the images were deconvolved using QuPath^50^ to unmix the primary and secondary stains, and background to three separate channels. Once GFAP, Aβ, and pTau-only images have been obtained, a final thresholding step individualized for each slice was taken to exclude non-specific staining and to allow for a subsequent % area calculation.

### Histology-MRI co-registration

The high-resolution MR images served as anatomical references for the registration of histological images. Regions in the histological images that exhibited significant divergence from the wet tissue state (i.e., MR images) due to deformation were manually excluded while preserving the image aspect ratio. Following the convergence of 2D affine co-registration of histology and MR images using the Image Processing Toolbox in MATLAB, a subsequent 2D diffeomorphic registration refinement was conducted between the histology image slices and MRI volumes. This refinement aimed to recover the true in-plane tissue shape and address remaining differences between the modalities. The diffeomorphic registration procedure utilized an efficient implementation of the greedy diffeomorphic algorithm,^51^ available as an open-source software package (greedy, https://github.com/pyushkevich/greedy). The greedy software was initialized and utilized as previously described.^52^ The transformed histology images were superimposed on MR images to evaluate the quality of co-registration. The study incorporated three sets of 29 tissue slices each, featuring GFAP, Aβ, and pTau immunostains, across 13 subjects.

### T_1_ and T_2_ maps and diffusion tensor MRI processing

Diffusion tensor imaging parameters,^53^ axial diffusivity (AD), radial diffusivity (RD), apparent diffusion coefficient (ADC), and fractional anisotropy (FA), were calculated using in-house MATLAB (The Mathworks, Natick, MA) code based on previous work.^54^

Conventional quantitative relaxation maps were first computed by fitting the signal decay to monoexponential functions. The T_1_ value was computed by fitting inversion recovery data that included 20 images with inversion times logarithmically sampled in the range of 14.6 ms and 980 ms, and echo time of 12.3 ms. The T_2_ value was computed by fitting a subset of the multidimensional data that included 20 images with echo times in the range of 12.3 ms and 125 ms.

We also applied a commonly used strategy to correct for possible between-subject differences arising from postmortem effects; we adjusted each voxel-averaged MRI parameter by dividing them by the mean for that parameter across GM voxels in each brain sample.

### Image domain masks and regions of interest

A *k*-means procedure with two clusters was used with the voxelwise MD-T_2_ distributions as inputs, to derive WM and GM image masks. Using image multiplication of the co-registered GFAP, Aβ, and pTau full resolution thresholded images and their inverted versions (i.e., binary complement) the following eight groups were generated according to neuropathology in the cortex: GFAP+/Aβ+/pTau+, GFAP-/Aβ-/pTau-, GFAP+/Aβ-/pTau-, GFAP+/Aβ-/pTau+, GFAP+/Aβ+/pTau-, GFAP-/Aβ+/pTau+, GFAP-/Aβ-/pTau+, and GFAP-/Aβ+/pTau-, where, for example, GFAP+/Aβ+/pTau+ is a mask containing only pixels that are positive for all immunostains.

Using the above masks, each image voxel was defined either as containing or not containing GFAP, Aβ, and pTau pathology. This process created eight groups of voxels across our study, from which 12 regions of interest (ROIs) from each group were selected (blind to MRI) for analysis, yielding a balanced histopathology representation. Despite being derived from the temporal pole, Case 5 (Table 1) was included in the study because it was one of the only samples containing GAP-/Abeta-/pTau+ ROIs, which are needed in order to avoid a confounded regression analysis. After extracting the ROIs, GFAP, Aβ, and pTau density were expressed as the percentage of total area within the ROI in the binary deconvolved images. In total, 96 regions of interest from 29 slices and 13 cases were included in this study.

### Multidimensional MRI processing

Before analysis, denoising using an adaptive nonlocal multispectral filter was performed on the MD-MRI data.^57^ The pre-processed data then underwent marginally constrained, ℓ_2_-regularized, nonnegative least square optimization to calculate the multidimensional distribution in each voxel, following established protocols.^42,58^ This approach, recognized for its robustness and reliability,^59–62^ resulted in T_2_-MD distributions for each voxel in this study.

We implemented the following procedure to correct for possible inter-site differences arising from postmortem effects: First, the GM mask was applied, and the maximal peak location in the spectral domain (i.e., T_2_-MD) was automatically found (this step was repeated for each subject). We arbitrarily set the JHU site as reference, to which UWA cases were aligned to in the spectral domain. This procedure ensures standardization across subjects, equivalent to the well-established strategy employed for voxel-average images.^55,56^

### Identification of abnormal T_2_-MD sub-voxel spectral component

Treating MD-MRI distributions as spectra raises the possibility to produce images of distinct spectral components through integration across a predefined parameter range, which can be thought of as a “spectral” ROI. The resulting integral value, ranging from 0 to 1, signifies a signal fraction within a particular multidimensional distribution. This computation can be applied voxel-wise to create an image highlighting that specific spectral component.^63^

While the T_2_-MD spectra contain rich information from multiple tissue components, we focused here on discovering spectral substrates that can distinguish cases with and without dementia. Based on our hypothesis that cortical astrogliosis is an important contributing factor to cognitive decline, we used the histologically derived information to search for differences between control, resilient and AD dementia cases in their respective T_2_-MD spectra. Specifically, cortical GFAP positive voxels (obtained using GFAP+ masks) were assumed to contain abnormal level of reactive astrocytes and were used to derive an average T_2_-MD spectral signature characteristic of astrogliosis. Applying the same principles, cortical GFAP negative voxels (obtained using GFAP-masks) were assumed to not contain any reactive astrocytes and were used to derive an average ‘normal’ T_2_-MD spectral signature. With the goal of identifying a spectral region that does not overlap, the two characteristic T_2_-MD spectral signatures, GFAP- and GFAP+, were then thresholded at 4% and 8% from maximal intensity, respectively. Finally, an astrogliosis-specific spectral ROI was obtained by taking the set difference of GFAP+ and GFAP-, which is the set of elements in the “astrogliosis” and not the “normal” T_2_-MD characteristic spectral signature. This process resulted in a spectral ROI, which was then applied voxelwise to create a map of astrogliosis specific spectral component.

### Statistical analysis

A multiple linear regression model was used to investigate associations of the MRI variables with astrogliosis, Aβ, and pTau pathologies due to AD. Prior to statistical analysis, all MRI markers were z-normalized (i.e., standardized).

We assessed effects of GFAP, Aβ, and pTau reactivity on MRI parameters, with each mean MRI metric within each ROI as the dependent variable. The model is given by: *P*_*i*_ = *β*_0_+ *β*_GFAP_\**GFAP* + *β*_Aβ_*Aβ + *β*_pTau_*p*Tau* +*β*_sex_\**sex* + *β*_age_\**age*, where *P*_*i*_is the mean ROI value of the parameter of interest (i.e., MD-MRI-derived astrogliosis marker, FA, AD, RD, T_1_, and T_2_) of the *i*th ROI. Sex and age were accounted for. Results are presented as the beta coefficients of estimates *β*_GFAP_, *β*_Aβ_, and *β*_pTau_, which due to standardization represents the standard deviation change in MR variable per % area staining of the respective pathology.

False discovery rate (FDR) correction was done to correct for multiple comparisons^64^ and the threshold for statistical significance was *p*_FDR_ < 0.05.

Multiple group analyses were performed using one-way ANOVA to determine differences in histopathological burden of different stains between AD dementia and resilient groups.

### Data availability

The datasets generated and analyzed during the current study are available from the corresponding author upon request.

## Results

### Cortical neuropathology

Table 1 summarizes the main demographic data, antemortem cognitive state (dementia vs. non-dementia), Braak stage, CERAD score, and region of the brain.

Figure 1A shows simultaneous views from cortical regions in three representative cases, control, resilient, and AD dementia, immunostained for markers of Aβ (4G8), pTau (AT8), reactive astrocytes (GFAP), microglia (IBA1), and phosphorylated transactive response DNA binding protein 43 (pTDP-43). Regional Aβ-plaque and pTau burden did not show apparent differences between the resilient and AD dementia cases, in agreement with previous observations.^16,22^ Higher burden of activated astrocytes can be seen in the AD dementia case compared to both control and resilient brains. Levels of microglia and pTDP-43 densities did not show any marked differences between subjects.

**Figure 1.**
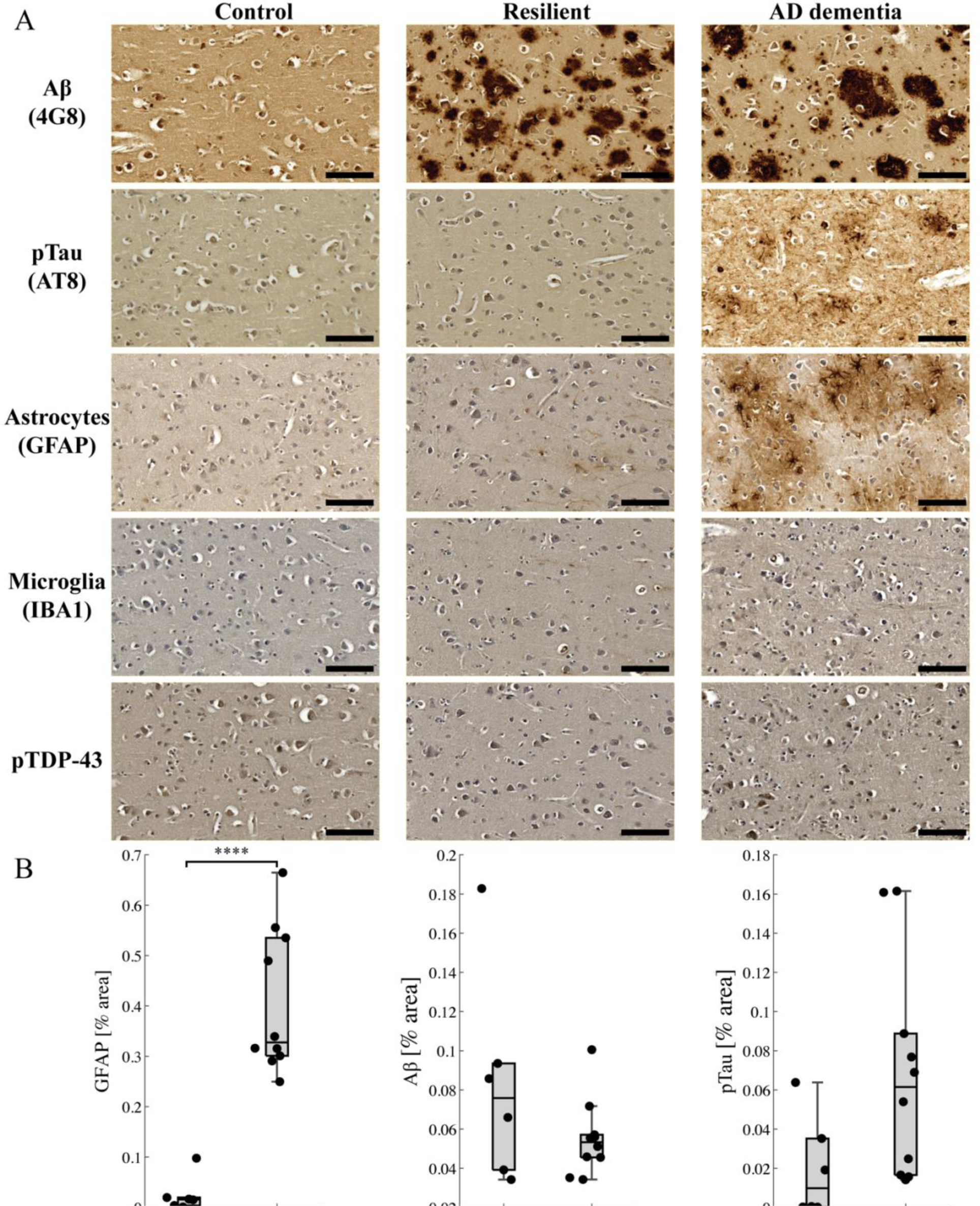
Histological findings in the cortex. **(A)** 4G8 (Aβ), AT8 (pTau), GFAP (astrocytes), IBA1 (microglia), and pTDP-43 sections from control, resilient and AD dementia representative cases containing superior frontal gyrus. Scale bar is 100 μm. **(B)** Quantifications for total GFAP, Aβ, and pTau burden (percent area) in superior frontal gyrus showed statistically significant differences between AD dementia and resilient brains for GFAP only, and not for Aβ or pTau.

After categorizing the AD dementia and resilient cases into two groups (two slices from each sample, total of 16), quantitative analysis was conducted to establish differences with respect to the Aβ, pTau, and GFAP burden in the cortex. The results, shown in Fig. 1B, showed statistically significant differences between AD dementia and resilient samples only for GFAP burden, and not for Aβ or pTau, indicating that the cognitive outcomes of these two groups were not predictable based on either regional Aβ-plaque or pTau loads.

Figure 2 and Supplementary Fig. 1 show overlayed deconvolved Aβ, pTau, and GFAP co-registered images of representative cases from the study, demonstrating pathological heterogeneity. Clear Aβ burden can be seen in all AD dementia and resilient cases. The presence of NFTs, however, is inconsistent, with an example of an AD dementia case that lacked pTau burden in the region (Fig. 2B), and an example of a control brain with low level pTau pathology (SI Fig. 1A). Astrogliosis as reflected by GFAP burden can be seen in the brain from all donors with dementia (Fig. 2A-B, SI Fig. 1B), and in none of the resilient and control cases.

**Figure 2.**
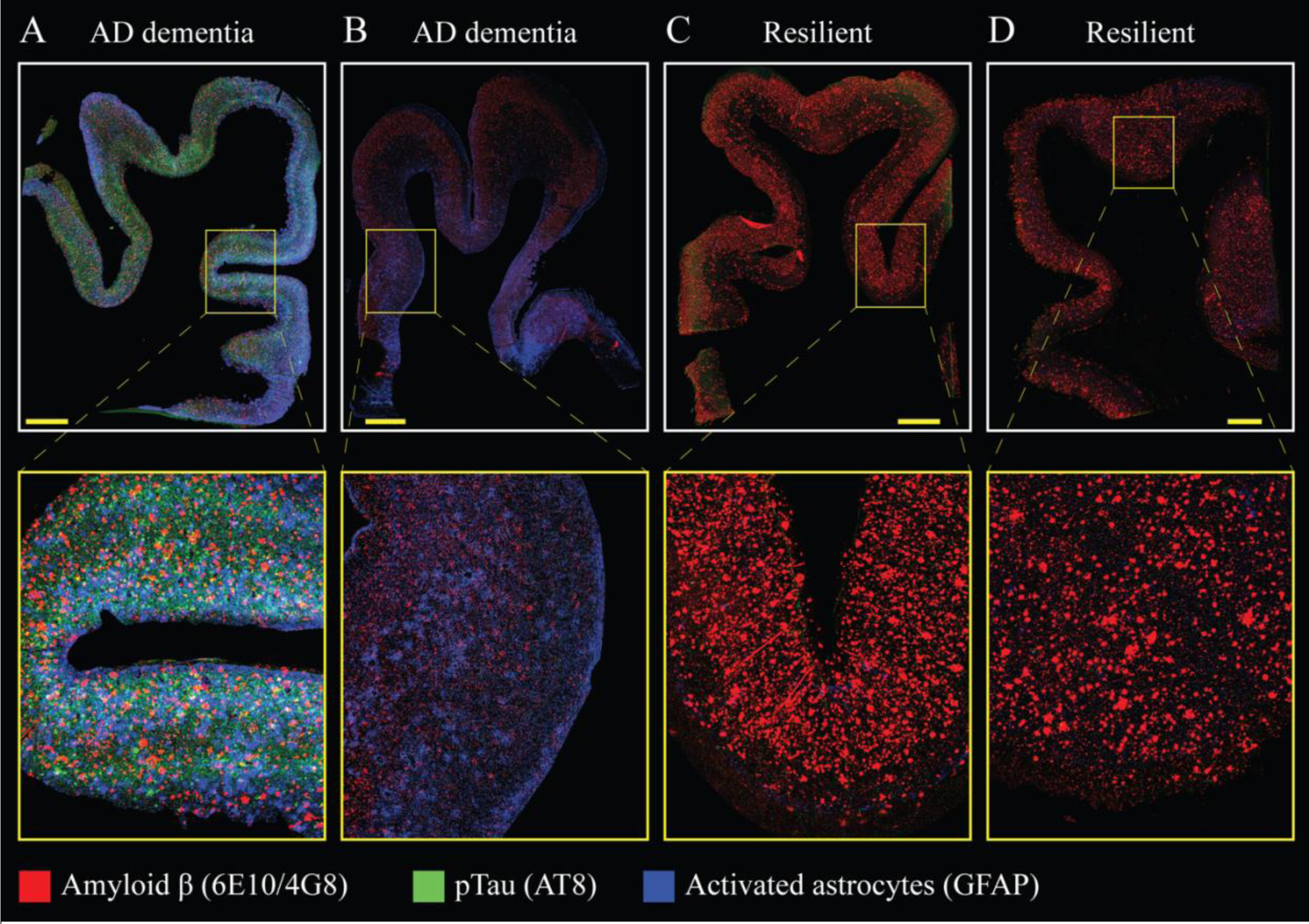
Whole sample visualization of co-registered immunohistochemistry sections. Immunostained Aβ, pTau, and GFAP images are co-registered and deconvolved, then overlayed as a red-green-blue image to illustrate the degree of pathological overlap in four representative AD dementia **(A-B)** and resilient **(C-D)** cases. Clear Aβ plaques can be seen in all cases (red), but only AD dementia cases exhibit abnormal GFAP burden. Scale bar is 2 mm.

### Cognitive state and astrogliosis modulate cortical microstructural properties

The carefully co-registered MRI and microscopy images facilitate quantitative multimodal analyses, and the high-resolution, high specificity microscopy data act as a pseudo ground truth estimates of tissue microstructure against which MR metrics can be compared. To assess the extent to which antemortem cognitive state is driving changes in the MR diffusion-relaxation response, we focused on reactive astrocytes and first produced characteristic T_2_-MD spectral signatures for GFAP negative cortical voxels from the control and resilient groups, shown in Figs. 3 A and B, respectively. Similarly, a characteristic T_2_-MD spectral signature for GFAP positive cortical voxels from the AD dementia group was generated and is shown in Fig. 3C. A visualization of these three characteristic spectral signatures together is shown in Fig. 3D. While the control and resilient signatures mostly overlap, the T_2_-MD characteristic distribution of the AD dementia group clearly diverges from the GFAP negative signatures. The diverging spectral region, illustrated in Fig. 3E, can be obtained by taking the set difference of GFAP+ and GFAP-, which is the set of elements in the AD dementia and not the control or the resilient T_2_-MD characteristic spectral signature. This process resulted in a spectral ROI (Fig. 3F), which was then applied voxel-wise to create images of abnormal T_2_-MD sub-voxel spectral component with expected specificity towards astrogliosis.

**Figure 3.**
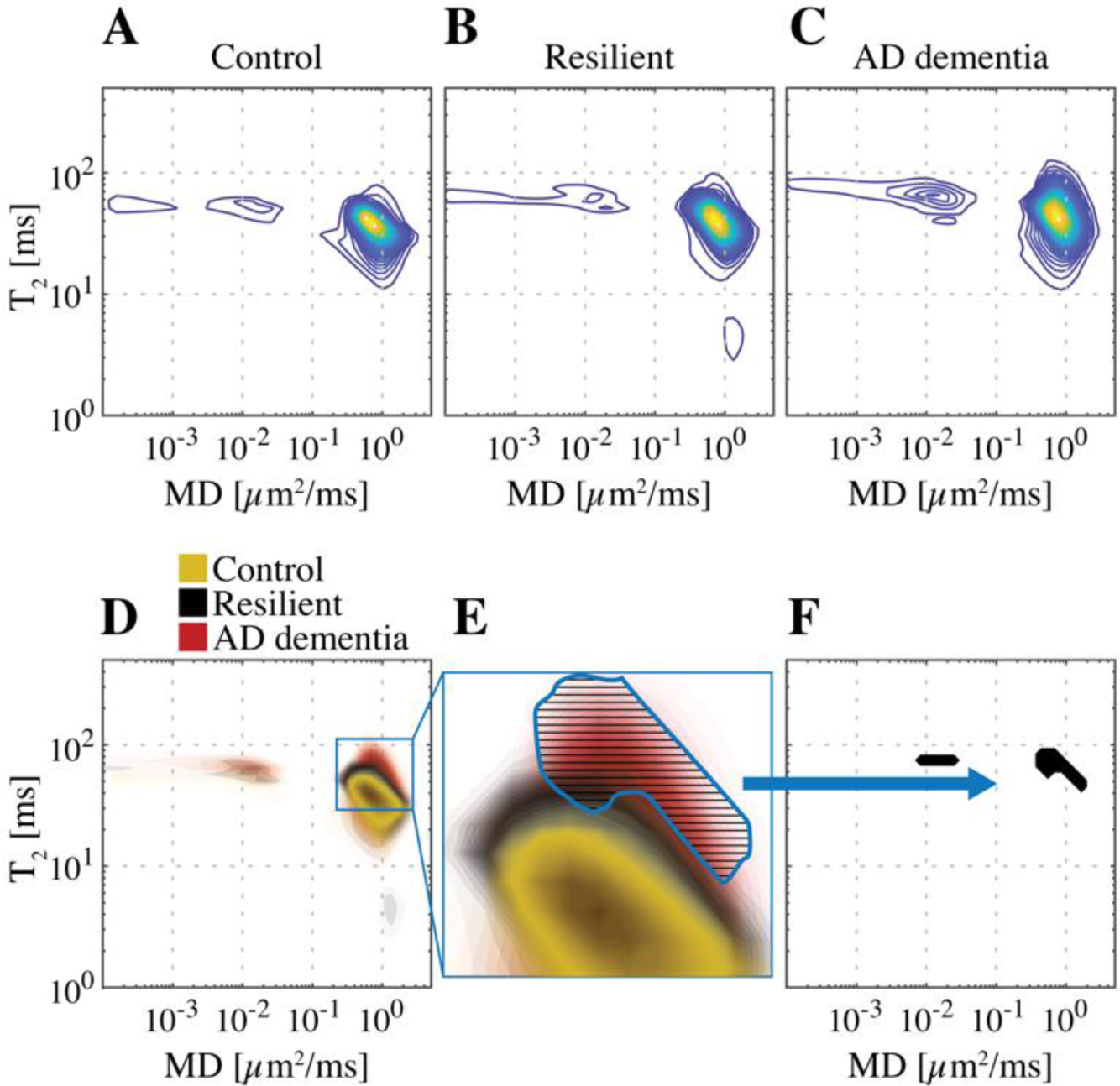
Cortical astrogliosis has a diffusion-relaxation spectral signature. T_2_-diffusion spectra in the cortex averaged across GFAP-negative voxels in **(A)** control and **(B)** resilient cases, and across GFAP-positive voxels in **(C)** AD dementia subjects. **(D)** A superposition of the average spectra from the three groups shows a unique spectral signature (marked in red), characteristic of the AD dementia cases, further magnified in **(E)**. Taking the set difference of GFAP-positive and GFAP-negative spectra results in **(F)** a spectral ROI, which can then be applied voxelwise to create a map of astrogliosis specific spectral component.

### Astrogliosis imaging in the cortex

Based on the control, resilient, and AD dementia cases we examined here, we have identified a T_2_-MD region in which most of the spectral information regarding astrogliosis pathology resides. Integration according to this spectral ROI, as detailed in the Materials and Methods section, was performed, resulting in MD-MRI astrogliosis biomarker maps.

Figure 4 shows conventional MRI and DTI maps, multidimensional MRI maps, and histological GFAP density images of six representative control, resilient, and AD dementia cases. Upon qualitative comparison, the MD-MRI neuropathology maps closely align with GFAP density images. While conventional MRI and DTI maps offer valuable macroscopic anatomical details such as gray-white matter separation, they fall short in capturing the microstructural or compositional alterations induced by astrogliosis, likely attributed to their inherent voxel averaging.

**Figure 4.**
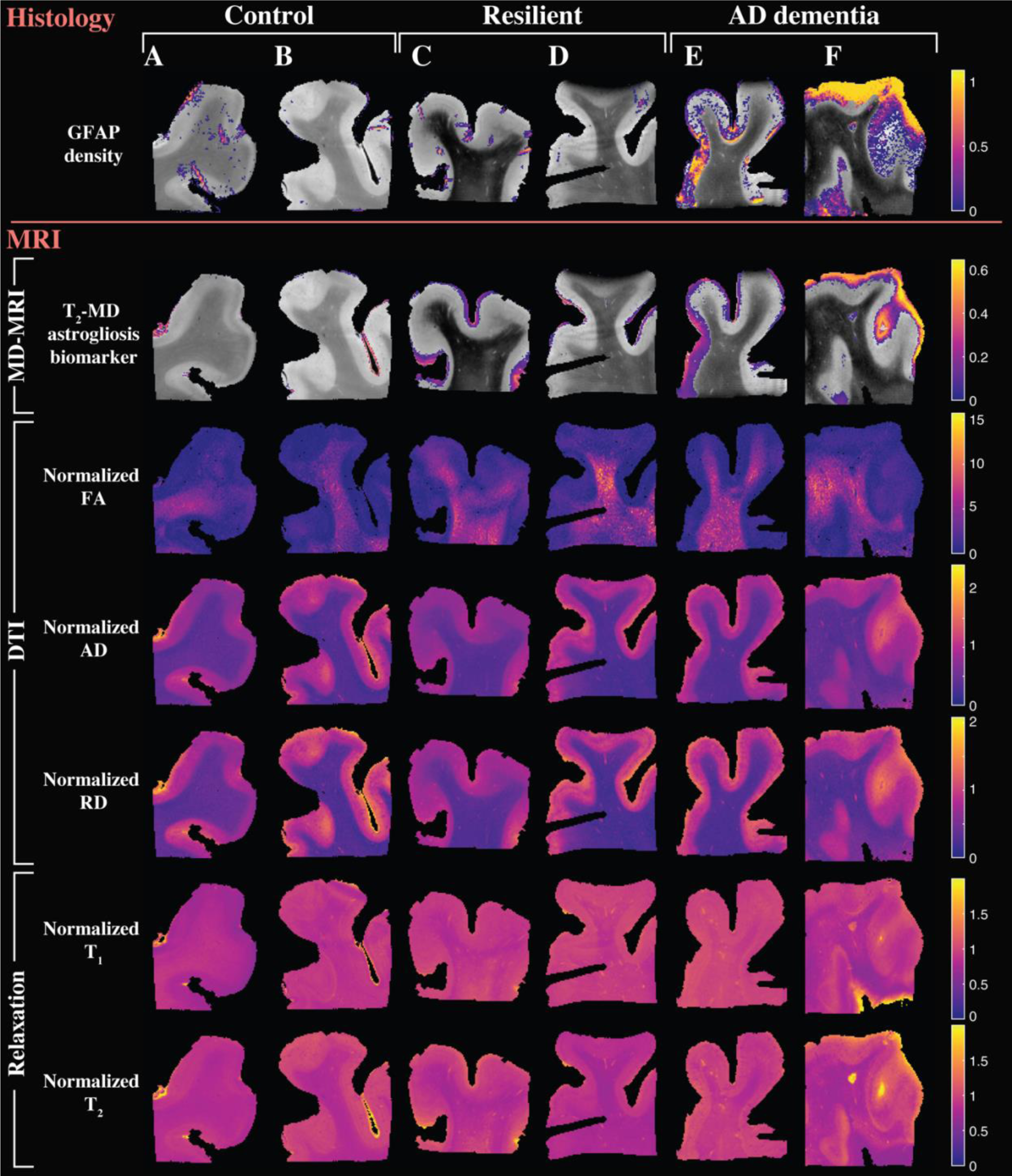
Multidimensional and voxel-averaged MRI maps. Control **(A-B)**, resilient **(C-D)**, and AD dementia **(E-F)** subjects are shown. The different rows correspond to the different MRI contrasts, including all the conventional relaxation and DTI parameters, and the proposed MD-MRI astrogliosis maps. In addition, the co-registered histological GFAP density maps are shown. To facilitate visualization, MD-MRI cortical astrogliosis maps were overlaid onto proton density images. These show substantial cortical pathology, while conventional relaxation and DTI maps do not show visible abnormalities.

### MD-MRI images astrogliosis and not Aβ or pTau pathology

The comprehensive multimodal dataset in this study, encompassing MRI volumes co-registered with GFAP, Aβ, and pTau histological slices, enables the investigation of the associations between the identified MD-MRI-derived biomarkers and their corresponding neuropathological drivers.

To validate our results, a robust image co-registration method was applied to accurately match MRI images and histopathology, after which each image voxel was defined either as containing or not containing GFAP, Aβ, and pTau pathology (e.g., [GFAP+/Aβ-/pTau-] is a voxel without any Aβ or pTau and with GFAP reactivity). This process created eight groups, from which 12 regions of interest (ROIs) were selected (blind to MRI) for analysis, reaching a total of 96 ROIs, with a balanced histopathology representation. We performed multiple linear regression to determine the relationship between MRI as the dependent variable, and GFAP, Aβ, and pTau % area staining levels as independent variables, adjusted for age and sex. Results are presented as the beta coefficients of estimates *β*_GFAP_, which due to standardization represent the standard deviation change in MR variable, per standard deviation change in GFAP % area. As shown in Fig. 5A, a strong and significant correlation was found only between the MD-MRI astrogliosis image intensity and GFAP immunoreactivity (standardized β_GFAP_ of 0.658, *p*_FDR_<0.0001). From the conventional voxel-averaged images, we found that GFAP density was insignificantly associated with FA (β_GFAP_=-0.251, *p*_FDR_=0.051), ADC (β_GFAP_=0.047, *p*_FDR_=0.589), AD (β_GFAP_=0.040, *p*_FDR_=0.634), RD (β_GFAP_=0.052, *p*_FDR_=0.586), T_1_ (β_GFAP_= −0.225, *p*_FDR_=0.059), and T_2_ (β_GFAP_= 0.200, *p*_FDR_=0.103), as displayed in Figs. 5B-G.

**Figure 5.**
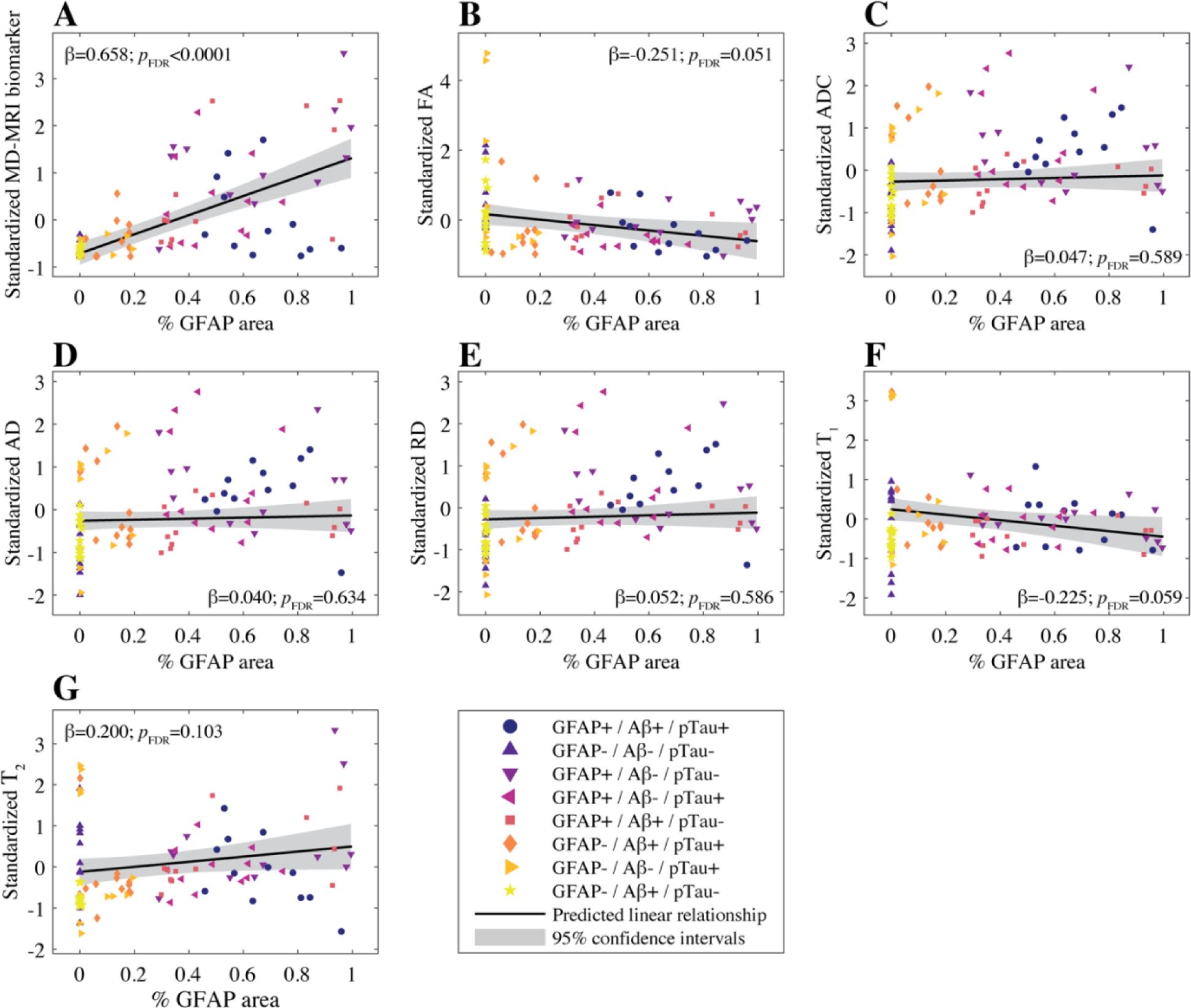
Radiological-pathological correlations between MRI metrics and GFAP density. GFAP density (% area) from 96 tissue regions from 13 subjects (color-coded, see legend) and the corresponding MR parameter correlations. Individual data points represent the mean value from each postmortem tissue sample. Scatterplots of the mean % GFAP area and **(A)** T_2_-MD cortical MD-MRI biomarkers show strong positive and significant correlation with GFAP density (standardized β_GFAP_ of 0.658, p_FDR_<0.0001). The conventional MRI metrics in **(B)**-**(G)** did not result in strong and significant correlations with % area GFAP.

The strong correlation between the MD-MRI-based neuropathological biomarker and GFAP indicates high sensitivity. The regression coefficients from the full model provided in Table 2 establish the specificity of this approach to astrogliosis, as evidenced by weak and statistically insignificant associations with Aβ and pTau immunostaining burden (β_Aβ_ = 0.009, *p*_FDR_=0.913 and β_pTau_ =-0.196, *p*_FDR_=0.051).

**Table 2.** Multiple linear regression analysis of the histological predictors of MD- and conventional MRI measures.

| Measure | Standardized $\beta$ ($p_{FDR}$-value) | | |
| --- | --- | --- | --- |
|  | GFAP | 6E10/4G8 | AT8 |
| MD-MRI biomarker | **0.658 (<0.001)** | 0.009 (0.913) | -0.196 (0.051) |
| FA | -0.251 (0.051) | 0.194 (0.103) | **-0.271 (0.033)** |
| ADC | 0.047 (0.589) | -0.155 (0.080) | **0.472 (<0.001)** |
| AD | 0.040 (0.634) | -0.139 (0.103) | **0.462 (<0.001)** |
| RD | 0.052 (0.586) | -0.163 (0.077) | **0.475 (<0.001)** |
| T<sub>1</sub> | -0.225 (0.059) | -0.198 (0.080) | **-0.462 (<0.001)** |
| T<sub>2</sub> | 0.200 (0.103) | -0.110 (0.392) | 0.118 (0.358) |

FDR-corrected $p$ -values are indicated in parentheses.
Multidimensional (MD); Fractional anisotropy (FA); Apparent diffusion coefficient (ADC); Axial diffusivity (AD); Radial diffusivity (RD); Glial fibrillary acidic protein (GFAP).

Although not the subject of this study, all DTI metrics and T_1_ were found to be significantly correlated with pTau immunoreactivity, exhibiting reduction in T_1_, as well as reduced FA and corresponding increases in diffusivity parameters, with increased pTau density burden (Table 2).

## Discussion

This study represents the first demonstration of noninvasive *ex vivo* differentiation between individuals with dementia and those with intact cognition, both exhibiting AD neuropathology. Prior studies and our own quantitative histological analysis found that astrogliosis induced GFAP burden distinguishes resilient versus AD dementia cases, suggesting cognitive outcomes could not be predicted based solely on Aβ or pTau loads. We therefore sought to show that in the presence of AD neuropathological changes, MD-MRI astrogliosis imaging in the human cortex can predict cognitive resilience. We demonstrated that cortical astrogliosis is associated with microstructural and compositional changes that result in a distinct *ex vivo* MD-MRI spectral signature, which can be leveraged to generate maps that quantify the severity of the astroglial neuropathology. Furthermore, through a multimodal approach integrating histology and MRI, we leverage the targeted specificity of immunostaining to demonstrate that cortical astrogliosis, rather than Aβ or pTau, underlies the pathological characteristic signature identified using MD-MRI.

An increasing body of research highlights the significant role of astrocyte reactivity in the pathophysiology of AD and other neurodegenerative disorders.^65,66^ The heterogeneous functional and morphological states of reactive astrocytes, along with their spatial distribution in the brain, are believed to reflect changes in brain homeostasis.^67^ In light of astrocytes’ importance, recent advancements in plasma GFAP assays have been introduced and evaluated in extensive imaging cohorts.^68^ Although evidence suggests a potential association between elevated plasma GFAP levels and Aβ pathology,^69,70^ recent findings indicate a strong negative correlation between cortical and plasma GFAP levels in both mice^71^ and humans.^72^ These discrepancies imply a complex relationship between plasma GFAP concentration, which is a global measure, and cortical astrocyte activation, likely influenced by peripheral tissues such as the liver or pancreas.^73^ Perhaps more importantly, the presence of astrogliosis in WM, which does not predict cognitive resilience, elevates global plasma GFAP levels, further weakening its link to cortical astrogliosis. These circumstances and limitations highlight the potential impact of an imaging modality for cortical astrogliosis mapping.

Multidimensional MRI allowed us to capture the strong association of T_2_ lengthening and diffusivity increase with the severity of abnormal astroglial reactivity in cortical GM regions. Increases in both relaxation times and diffusivities point to a reduction in the degree of neuronal density that is expected to result from the presence of highly reactive astrocytes,^74,75^ as illustrated in Fig. 6. Similar relaxation-diffusion correlation trends were recently observed and reported in astrogliosis in WM.^42^ However, while the presence of reactive astrocytes in cortical GM and WM modulates their respective characteristic MD-MRI signatures in similar ways, the affected T_2_ and diffusivity values are markedly different. The differences in T_2_ and diffusivity values within WM and GM are well-documented, and are driven by decreased myelin content, increased cellular size, decreased anisotropy, and reduced orientational coherence in GM compared to WM.^76–79^ The unique ability of MD-MRI to access specific intra-voxel diffusion-relaxation regions allows one to exploit this inherent WM-GM contrast, and separately image GM and WM astrogliosis.

**Figure 6.**
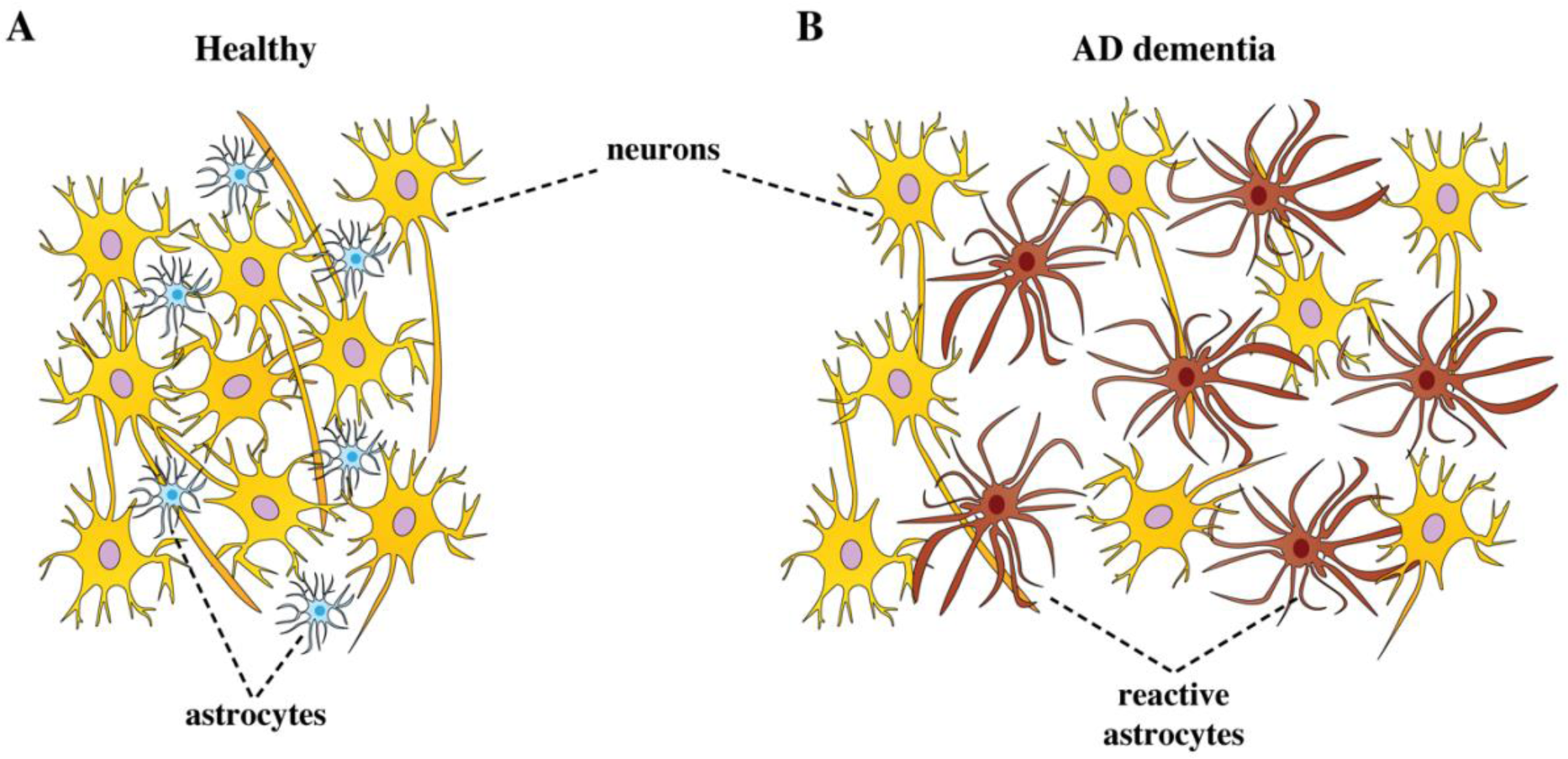
Illustration of microstructural changes occurring in the cortex when astrogliosis is present. In **(A)** neurons are tightly aligned forming a densely packed cellular environment. In **(B)** reactive astrocytes display hypertrophy and have overlapping processes that disrupt the tissue microstructure. These changes are hypothesized to be reducing the overall cellular density in the gray matter cortical regions, rendering them detectable via MD-MRI.

Because MRI signals are often indirect measures of the brain features of interest, determining the microstructural underpinnings of a given MRI change is impossible without obtaining complementary and independent information. To achieve this specificity, our approach in this study spans modalities (*ex vivo* MRI and microscopy) and orders of spatial magnitude. Crucially, the MRI and microscopy images were carefully co-registered together to facilitate quantitative multimodal analyses, and the high-resolution, high specificity microscopy data act as a pseudo ground truth estimate of tissue microstructure against which MR metrics can be compared. Including Aβ, pTau, and GFAP cortical microscopy and their corresponding staining burdens allowed to test their associations with the MD-MRI signature we discovered (Fig. 3), and further, to pinpoint to what degree each of these neuropathologies drive the MR signal change. Our results showed that it is astroglial reactivity, and not Aβ or pTau, that is driving the microstructural changes that are reflected in the MD-MRI signature, as evident in Table 2 (β_GFAP_=0.658, *p*_FDR_<0.0001). These findings represent specificity towards astrogliosis in the presence of Aβ and pTau pathologies.

When used conventionally, diffusion and relaxation are encoded separately, i.e., in a one-dimensional (1D) manner. A wide range of 1D diffusion MRI models, including DTI and multicomponent biophysical models,^80,81^ provide excellent sensitivity to microstructural changes in WM. However, their limitations in studying GM due to deviations from assumed conditions are well-documented.^78,82^ These limitations explain the weak associations between the various 1D conventional MRI (DTI and relaxometry) and the presence of cortical astrogliosis (Fig. 5 and Table 2). The primary reason for this lack of sensitivity is that these methods do not allow for the selective extraction of the pathology-affected spectral range per subject, as they provide only scalar averages, rather than distributions. In addition, we showed here and previously^42^ that if used separately, any 1D relaxation or diffusion MRI measurement lacks the sensitivity to disentangle the microstructural alterations induced by astrogliosis. On the other hand, we found that increased pTau pathology is strongly correlated with DTI parameters and with T_1_ (Table 2). The positive correlations between DTI diffusivities and pTau staining burden, along with the negative correlations with FA and T_1_, can be explained by the disruption of the cortical laminar structure^83^ caused by tauopathy. Echoing these results, findings from *ex vivo* mice and human studies showed significant correlations of pTau with diffusion MRI metrics in cortical GM.^84,85^ Put together, previous and current results encourage the future investigation of a T_1_-diffusion MD-MRI framework focused on directly mapping pTau pathology in the cortex.

A limitation of our study is a difference in neurofibrillary tangle severity, as denoted by Braak staging between the AD dementia cases and resilient cases; therefore, the resilient cases may in fact represent the pre-clinical stage of disease. However, for all cases, the degree of pathology present is considered sufficient to cause dementia; further, in the tissue sections examined for this study, the total pTau burden was not significantly different between the AD dementia and resilient cases. In addition, like all *ex vivo* human MRI studies, our data are influenced by postmortem factors like degeneration and fixation-induced dehydration. This fixation process itself alters tissue properties, precluding direct comparison with *in vivo* data.^86,87^ Here, postmortem MRI functions as a crucial intermediary between histology and *in vivo* imaging, filling the knowledge gaps in understanding MRI signal response to AD pathology. Although we successfully showcased the capability to directly visualize the aberrant neuroinflammatory response to AD by accessing sub-voxel components using MD-MRI in *ex vivo* conditions, translating these measurements to humans requires noninvasive imaging techniques applicable under *in vivo* conditions. In doing that, the challenges we face are mainly due to diffusion gradients hardware limitations, and limited scan time, compared with *ex vivo* work. Recent major advances in clinical translation^34,36^ led to implementations of diffusion-relaxation MD-MRI imaging protocols on clinical scanners with feasible scan times.^88,89^ Despite these *in vivo* MD-MRI protocols, differences remain between clinical and preclinical MD-MRI methodology, and therefore the current *ex vivo* MD-MRI cortical astrogliosis imaging findings remain to be demonstrated *in vivo*.

While a clear link between glial activation in AD and cognitive impairment is emerging, the debate whether reactive astrocytes in the cortex directly contribute to neuronal damage and cognitive decline in AD, or simply reflect a response to injury, or both, is unresolved. Regardless, a noninvasive imaging method to visualize these glial changes holds promise as a predictor of cognitive state. Although specific *in vivo* detection of astroglial pathology remains a critical unmet need, we demonstrate here an MD-MRI signature of cortical astrogliosis in AD, translating into biomarker images that closely reflect GFAP pathology. As numerous potential disease-modifying therapies for AD advance to late-stage clinical trials^90,91^ and the absence of a definitive marker of neurodegeneration persists, MD-MRI astrogliosis imaging could play a crucial role as a potential indicator for monitoring the early stages of AD progression, for evaluating the response to anti-Aβ treatments in clinical trials, and, more significantly, for exploring new neuroinflammatory therapeutical avenues.

## Acknowledgments

We thank Dr. Rafa deCabo, Ms. Sara Abdo, and Ms. Ericka Oglesby for facilitating and assisting with scanning microscopy slides. We also thank Ms. Aimnee Schantz and Mr. John Campos for outstanding administrative support, and the brain donors and their families, without whom this research would be impossible. We would like to thank all the BLSA participants and families that contributed to brain donation.

## Abbreviations

AD: Alzheimer’s disease
GFAP: glial fibrillary acidic protein
ROI: region of interest
MD-MRI: multidimensional MRI
Aβ: amyloid β
NFTs: neurofibrillary tangles
pTau: hyperphosphorylated tau

