## Supplementary Fig for "Resiliency to Alzheimer’s disease neuropathology can be distinguished from dementia using cortical astrogliosis imaging"

### Supplementary Results

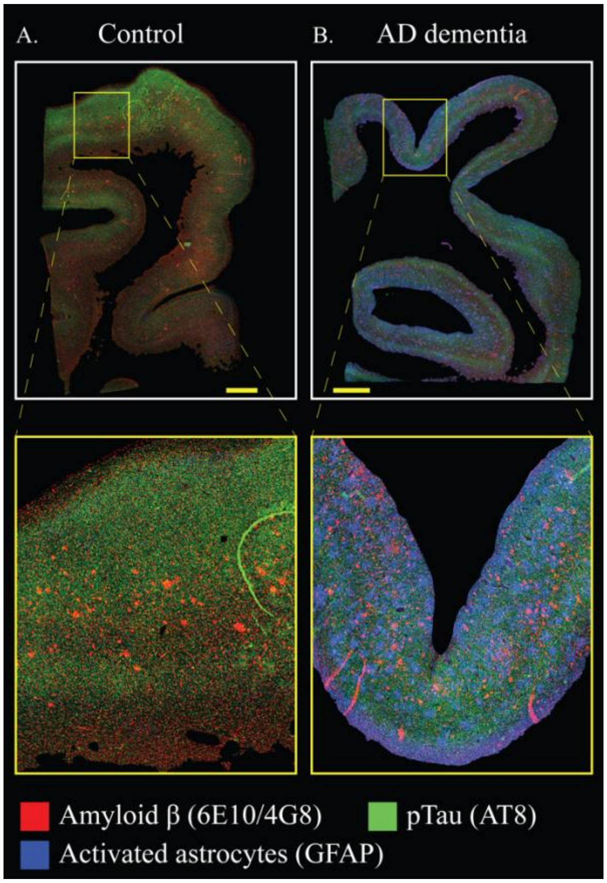


***Supplementary Figure 1. Whole sample visualization of co-registered immunohistochemistry sections.*** *Immunostained Aβ, pTau, and GFAP images are co-registered and deconvolved, then overlayed as a red-green-blue image to illustrate the degree of pathological overlap in a* ***(A)*** *representative control case with low tau pathology, and* ***(B)*** *an AD dementia case. Scale bar is 2 mm.*

### Supplementary Methods

#### MRI acquisition

The acquisition of multidimensional data was done using echo planar imaging (EPI) readout according to the MADCO framework encoding scheme,[1,2] and by varying the following two experimental parameters: the echo time, TE, and the diffusion weighting, *b*, providing $T_2$-, and diffusion-weighting, respectively.

The minimal TE value depends on the sample physical dimensions because of the varying imaging matrix size that is intended to keep the spatial resolution constant at 200 µm isotropic voxels. We kept the minimal TE relatively constant across the samples at 12.3±0.8 ms, by adjusting the number of EPI segments as necessary (<13).

The two 1D distributions of $T_2$ and MD were estimated, respectively, with the following data acquisition protocols: For $T_2$ encoding, a 1D $T_2$-weighted data set ($b$=0) with 20 logarithmically sampled TE values ranging from 12.3 to 125 ms by using a DWI-EPI sequence. For diffusion encoding, we used the isotropic generalized diffusion tensor MRI (IGDTI) acquisition protocol to achieve an efficient orientationally averaged DW signal[3] with the following parameters: 16 linearly sampled $b$-values ranging from 2,540 to 14,700 s/mm$^2$ in 3 directions, 14 linearly sampled $b$-values ranging from 4,140 to 14,700 s/mm$^2$ in 4 directions, and 9 linearly sampled $b$-values ranging from 8,260 to 14,700 s/mm$^2$ in 6 directions, using the efficient gradient sampling schemes in Table 2 in Avram *et al.*[3] This type of diffusion encoding increases the contrast given by local anisotropy and is not intended to measure the isotropic diffusion in the system. Additional diffusion parameters were gradient duration of $\delta$=4 ms and diffusion time of $\Delta$=15 ms.

The 2D distribution of MD-$T_2$ was estimated (in conjunction with the *a priori* obtained 1D distributions as constraints) from a 2D D-$T_2$-weighted data set with 16 sampled combinations of echo times and $b$-values within the aforementioned 1D acquisition range.

The data were averaged 4 times to maintain high signal-to-noise ratio (SNR), which was always maintained above 100 (defined as the ratio between the average unattenuated signal intensity within a tissue region of interest, and the standard deviation of the signal intensity within the background). The sample temperature was set at 16.8°C.

### Immunohistochemistry

University of Washington: Four µm-thick tissue sections were cut on a microtome and using previously optimized conditions, immunohistochemistry was performed using a Leica Bond Rx and Max Fully Automated IHC and ISH Staining Systems (Leica Biosystems, Wetzlar, Germany). Serial sections were immunostained with mouse monoclonal antibody against paired helical filament tau (AT8, 1:1,000 dilution) (Invitrogen, Carlsbad, CA), mouse monoclonal against β-amyloid (6E10, 1:1,000) (Bioledgend), rat monoclonal against phosphorylated TDP-43 (ser409/ser410, 1:1,000) (Millipore, Burlington, MA), rabbit polyclonal against IBA-1 (1:1,000) (Wako, Richmond, VA), and rabbit polyclonal against glial fibrillary acidic protein (GFAP) (1:1,000) (Dako, Santa Clara, CA). Appropriate positive and negative controls were included with each antibody and each run. Additionally, Nissl stains were performed according to accepted protocols.

Johns Hopkins University: Ten µm-thick tissue sections were cut on a microtome and using previously optimized conditions. Antigen retrieval and tissue re-hydration was then completed in the Lab Vision™ PT module (Fisher Scientific, Waltham, MA) using Epredia™ Dewax and HIER Buffer M (Fisher Scientific, Waltham, MA). Immunohistochemistry was completed using a Lab Vision™ Autostainer 360-2D, (Fisher Scientific, Waltham, MA). Serial sections were immunostained with mouse monoclonal antibody against paired helical filament tau (AT8, 1:200

dilution) (Thermo Fisher, Waltham, MA), mouse monoclonal against β-amyloid (4G8, 1:500) (BioLegend, San Diego, CA), rat monoclonal against phosphorylated TDP-43 (ser409/ser410, 1:200) (BioLegend, San Diego, CA), rabbit polyclonal against IBA-1 (1:500) (Wako, Richmond, VA), and chicken polyclonal against glial fibrillary acidic protein (GFAP) (1:500) (Abcam, Boston, MA). Appropriate positive and negative controls were included with each antibody and each run. Additionally, Nissl stains were performed according to accepted protocols.
